# Rapid delivery of RNase H1 enables replication fork progression under stress independently of R-loop resolution

**DOI:** 10.1101/2025.02.27.640516

**Authors:** Amandine Basille, Yea-Lih Lin, Connor Kirke, David Cluet, Yara Bawadi, Laetitia Vachez, Virgile Dervieux, Laura Guiguettaz, Pascal Bernard, Stephan Hamperl, Philippe Pasero, Emiliano P. Ricci, Vincent Vanoosthuyse

## Abstract

The over-expression of RNase H1 enables the progression of replication forks under stress and safeguards genome stability. The most common interpretation of these observations is that RNase H1 over-expression removes R-loops, whose formation would hinder replication fork progression. Here we challenge this model using an innovative strategy to rapidly deliver ready-made *Escherichia coli* RNase H1 (RnhA) in live cells. We show that RnhA is enriched in the vicinity of active replication forks and that it rescues replication fork progression in different stress conditions. However, RnhA had no impact on R-loops mapped by the DRIP or Cut&Tag methods. Our results reveal the existence of distinct populations of RNA:DNA hybrids that are differently sensitive to RnhA and invites new interpretations of well-established observations.

**HIGHLIGHTS:**

- Rapid delivery of R-loop sensors and regulators in live cells
- Rapid delivery of RnhA rescues replication fork progression in stress conditions
- Rapid delivery of RnhA does not impact R-loops
- DRIP detects RNA:DNA species insensitive to RNase H1 in cells

## INTRODUCTION

In differentiated cells, early DNA replication forks progress slower and pause more frequently than late replication forks. Strikingly, inhibition of gene transcription speeds up early forks, suggesting that active transcription in early S phase hinders fork progression ^1,2^. A large body of evidence now supports the idea that conflicts between transcription and replication are a threat to chromosome integrity and that cells have evolved a number of mechanisms to either prevent or mitigate the consequences of such Transcription-Replication Conflicts (TRCs) (reviewed in ^3^). Interestingly, TRCs might be more frequent in some tumours and to target the molecular response to TRCs could constitute a promising therapeutic strategy ^4–7^.

The reasons why active transcription threatens advancing replication forks are likely multifactorial. Both transcription and replication alter DNA topology, and excessive positive topological stress could mechanically impede the movement of replication forks by impairing DNA unwinding, particularly in the case of head-on conflicts between transcription and replication (reviewed in ^8^). Altered DNA topology also increases the likelihood of co-transcriptional R-loops, three-stranded chromatin structures that form when the nascent RNA winds around its DNA template resulting in a RNA:DNA hybrid and an unpaired non-template DNA strand ^9,10^. In support of the idea that excessive R-loop formation might impede the movement of replication forks and compromise genome integrity, a large body of evidence consistently shows that the adverse consequences of TRCs can be largely alleviated by over-expressing RNase H1, an enzyme that degrades the RNA moiety of RNA:DNA hybrids (reviewed in ^11^). Conversely, loss of RNase H1 results in the complete collapse of DNA replication at a single head-on TRC engineered in the genome of *Bacillus subtilis* ^12^.

While these concordant observations are very compelling in establishing that RNA:DNA hybrids could interfere with faithful DNA replication and the resolution of TRCs, they do not prove beyond doubt that toxic RNA:DNA hybrids originate from pre-existing, co-transcriptional R-loops. Recently, post-replicative RNA:DNA hybrids embedded in nascent DNA strands have been postulated to form behind active replication forks in response to TRCs and to interfere with post-replicative processes important for fork stability and restart ^13–16^. Whether these post-replicative RNA:DNA hybrids are by-products of pre-existing R-loops that have been transferred behind forks by stress-induced DNA transactions or whether they are generated *de novo* in response to replication stress remains poorly understood and could depend on the type of stress and the resulting fork configurations. To better understand how RNA:DNA metabolism impacts replication fork progression and stability, the source of toxic hybrids in different experimental situations must be formally identified. Henceforth, we will refer collectively to RNA:DNA hybrids that impact fork stability and restart under stress as DRIFs (for <u>D</u>NA:<u>R</u>NA hybrids with Impact on <u>F</u>orks).

To understand the impact of DRIFs on fork transactions, dedicated tools are needed. Currently, the most popular tool to manipulate RNA:DNA levels in live cells is the lengthy over-expression of RNase H1 in asynchronous cell populations. As R-loops have been proposed to be important transcription regulators (reviewed in ^17^), the long-term over-expression of RNase H1 could significantly alter the transcriptome in a way that only indirectly mitigates the consequences of TRCs (reviewed in ^18^). The lengthy over-expression of RNase H1 therefore appears ill-adapted to the study of DRIFs.

Here we built an alternative strategy to manipulate DRIF levels in live human cells. We used Virus-Like Particles (VLPs) to deliver ready-made *Escherichia coli* RNase H1 (RnhA) for only short periods of time, which did not substantially impact the transcriptome. We show that the rapid delivery of ready-made RnhA mitigates the impact of replication stress on replication fork progression but has no impact on R-loops mapped by DRIP or Cut&Tag. This strongly suggests that increased RNase H1 activity facilitates the progression of replication forks under stress independently of R-loop resolution, at least as R-loops are often defined and detected.

## RESULTS

### Rapid and flexible delivery of ready-made RNase H1 in the nuclei of live cells

Virus-Like Particles (VLPs) have previously been used to deliver ready-made genome editors such as Cas9 in live vertebrate cells (see for example ^19,20^). VLPs are produced by the multimerization of the retroviral Gag polyprotein at the plasma membrane, which induces the curvature of the membrane required for vesicle formation and subsequent budding. The resulting particles lack genetic information but inclusion of fusogenic viral envelope proteins allows them to fuse with target cells and to deliver their protein cargo (Fig. 1A). The fusion of a protein of interest to the C-terminus of Gag results in its inclusion in VLPs. The addition of a proteolytic site for the protease of the Gag polyprotein at the junction between Gag and the protein of interest ensures its delivery into cells as a separate entity (Fig. 1A). We produced VLPs carrying the RNase H1 enzyme from *Escherichia coli* (RnhA) fused at the N-terminus to the V5 epitope and at the C-terminus to a Nuclear Localisation Signal (NLS). The choice of RnhA was motivated by the fact that it has a high catalytic activity and that it is unlikely to be targeted by inactivating post-translational modifications or to interact with endogenous protein partners. The amount of RnhA in VLPs can be adjusted during production for dose-dependent delivery and two proteins can be loaded and delivered at the same time, establishing VLPs as a flexible protein delivery system (Fig. S1A). Immunostaining showed that VLP-delivered RnhA accumulated exponentially in nuclei in the first three hours of incubation before reaching a plateau after four hours (Fig. 1B). In nuclei, RnhA accumulated in the nucleolus and was also uniformly distributed over chromatin, in a manner reminiscent of the endogenous RNase H1 enzyme ^21^ (Fig. 1C). Fractionation of protein extracts established that ∼60% of VLP-delivered RnhA remained in the soluble fraction whilst ∼40% associated in a salt-sensitive manner with chromatin (Fig. S1B). To assess whether VLP-delivered RnhA could recognise and associate with RNA:DNA hybrids in cells, we delivered the D10N mutant of RnhA (RnhA^D10N^, thereafter called *d*RnhA) that can recognise but not degrade RNA:DNA hybrids ^22^. After immunoprecipitation of *d*RnhA from the chromatin-containing fraction, its associated nucleic acids were purified and analysed by slot blot for the presence of dsDNA and RNA:DNA hybrids. RNA:DNA hybrids were specifically enriched upon immunoprecipitation of *d*RnhA, confirming that VLP-delivered RnhA associates with RNA:DNA hybrids in human cell extracts (Fig. 1D). Taken together, these data show that VLPs can rapidly deliver different isoforms of RnhA in the nuclei of live human cells. As a comparison, it was necessary to wait ∼20 hours in HeLa cells to obtain a similar amount of human RNase H1 by transfection of an over-expression plasmid (Fig. 1E).

**Figure 1:**
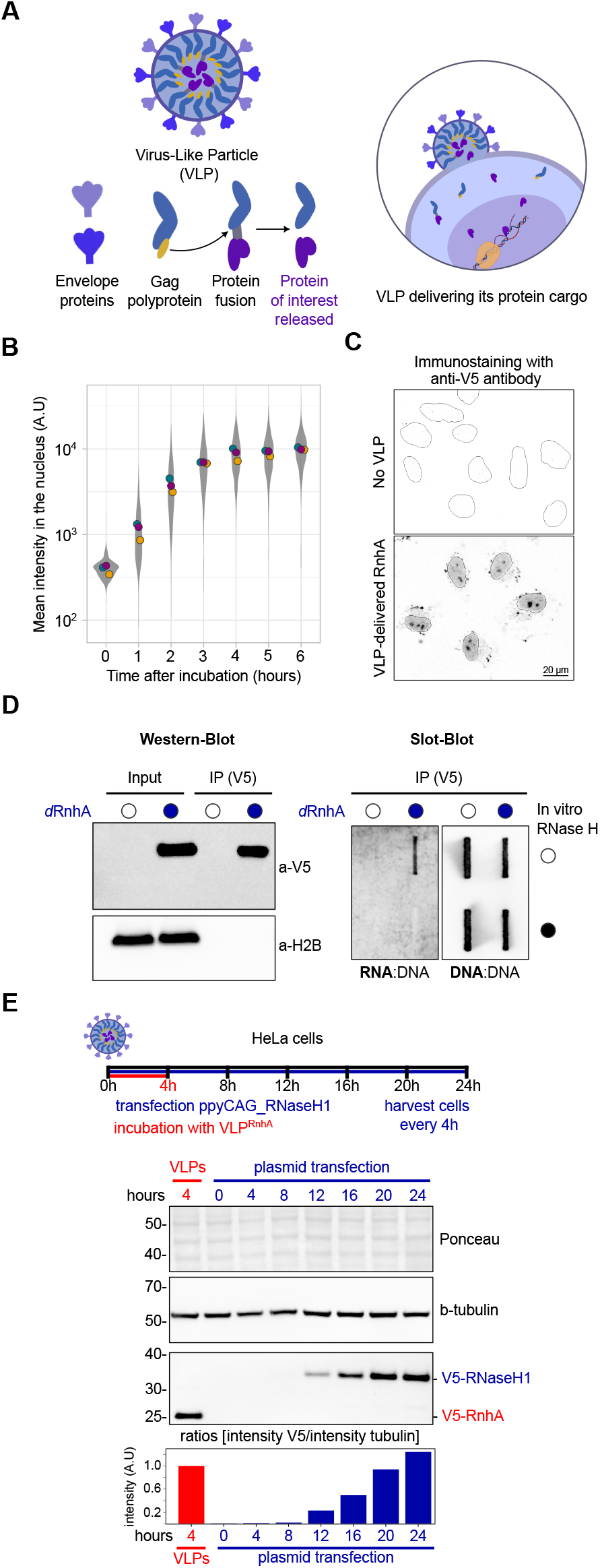
Virus-Like Particles (VLPs) rapidly deliver RnhA in the nuclei of live cells. A. Schematic representation of VLPs. The protease domain of the Gag polyprotein (in yellow) cleaves the junction between Gag and the protein of interest. As a result, VLPs deliver the ready-made protein of interest to target cells as a separate entity. **B**. Immuno-localisation was used to quantify the nuclear levels of VLP-delivered RnhA over time. **C**. Immuno-localisation of VLP-delivered RnhA at t = 4h. Cells not incubated with VLPs were used as controls. Nuclei borders are indicated by a solid line. **D**. Chromatin-associated, VLP-delivered *d*RnhA was immuno-precipitated (left) and its associated nuclei acids analysed by slot blot (right). **E**. Representative western blot analysis of the accumulation of V5-tagged, human RNase H1 over time upon transfection of an over-expression plasmid. The amount of V5-tagged RnhA delivered by VLPs after four hours is shown for comparison. Tubulin was used as a loading control. V5/tubulin ratios normalised over the VLP condition are shown below.

### VLP-delivered RnhA has a negligible impact on the steady state transcriptome

To evaluate the impact of VLP-delivered RnhA on the transcriptome, we carried out total RNA-seq analysis of HeLa cells treated with RnhA-containing VLPs for 4h or 24h (Fig. S2A). We used VLPs carrying GFP or *d*RnhA as controls. We detected only negligible changes to the transcriptome upon addition of VLPs delivering either control proteins or active RnhA, even after a 24-hour incubation (Fig. S2BC). These observations were confirmed when sequencing chromatin-associated RNAs (Fig. S2D). In sharp contrast, re-analysis of published RNA-seq data ^23^ showed comprehensive transcriptome perturbations upon over-expression of RNase H1 for 36 hours (Fig. S2E). In particular, transcription of the ubiquitin-like *ISG15* was strongly activated (Fig. S2F). As ISG15 conjugation to fork-associated proteins was previously shown to mitigate the consequences of replication stress ^24–26^, the sole activation of ISG15 might be sufficient to explain why RNase H1 over-expression mitigates the consequences of TRCs in those cells. To conclude, contrary to the lengthy over-expression of RNase H1, there are no significant confounding effects on the steady state transcriptome associated with the VLP-mediated delivery of either control proteins or active RnhA.

### VLP-delivered RnhA localises in the vicinity of replication forks

As the over-expression of RNase H1 has been shown to modulate the progression of replication forks in response to replication stress, we first asked whether VLP-delivered RnhA could be found in the vicinity of replication forks. We synchronised HeLa cells in S-phase using a single thymidine block (Fig. 2A) and used Proximity Ligation Assay (PLA) to measure the proximity between VLP-delivered proteins and the replisome component Proliferating Cell Nuclear Antigen (PCNA). As the different VLP-delivered proteins carry the same V5 tag in their N-terminus and the same NLS in their C-terminus, PLA signals for the different VLP-delivered proteins are directly comparable. We first confirmed that the delivery of the different proteins did not impact the quality of the synchronisation in S phase (Fig. 2B). Strikingly, only the delivery of the inactive *d*RnhA yielded strong PLA signals with PCNA (Fig. 2C). Immuno-staining experiments found no major differences in the levels of RnhA or *d*RnhA on chromatin (Fig. 2D), suggesting that the failure by RnhA to yield robust PLA signals is more likely explained by a short residence time on chromatin than by its depletion from chromatin. To quantify PLA signals in an unbiased manner, we used a machine-learning approach to classify nuclei into four different clusters depending on the strength of their signals (see Methods) (Fig. 2E). Interestingly, PLA signals with *d*RnhA were not significantly altered by a two-hour triptolide treatment (TPL, Fig. 2F), which efficiently depleted RNA polymerase II (RNAP2) from chromatin (Fig. 2G). This demonstrates that *d*RnhA does not require ongoing RNAP2 transcription to be positioned in the vicinity of PCNA. As *d*RnhA is unlikely to harbour determinants for targeting to specific genomic regions in human cells, its proximity to PCNA most likely reflects an association with transcription-independent DRIFs.

**Figure 2:**
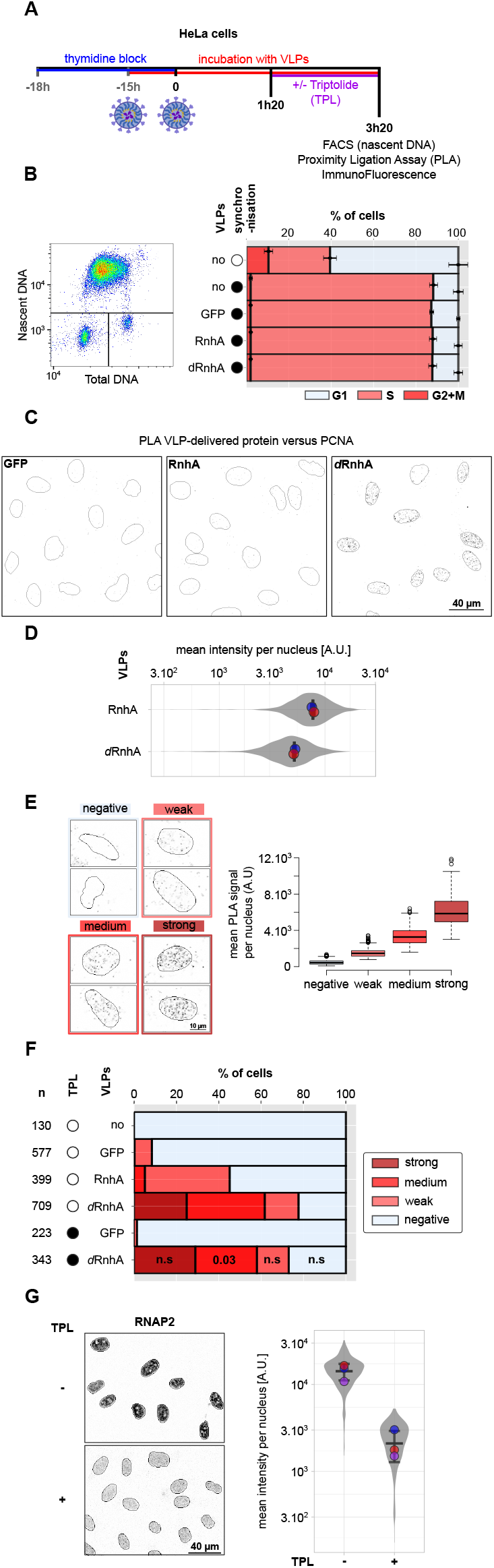
VLP-delivered RnhA is positioned in the vicinity of active replication forks. A. Schematic representation of the experiment. **B**. Cell-cycle profile of cells synchronised in S-phase and incubated with VLPs delivering the indicated proteins. (left) example of a typical FACS profile. (right) % of cells in the different cell-cycle phases. **C**. Examples of PLA signals in the indicated conditions. Nuclei borders are indicated by a solid line. **D**. Immuno-localisation was used to quantify the nuclear levels of VLP-delivered RnhA and *d*RnhA. **E**. (left) Representative nuclei for the different clusters. Nuclei borders are indicated by a solid line. (right) mean PLA signals per nucleus in the different clusters. **F**. % of nuclei in the different clusters per conditions. The total number of nuclei analysed over 3 to 5 biological replicates is indicated on the left (n). Similarity between *d*RnhA and *d*RnhA + TPL was tested using the Wilcoxon-Mann-Whitney test and the resulting p-values are indicated. n.s: not significant. **G**. The efficiency of the TPL treatment was validated by measuring the amount of RNAP2 on chromatin using Immuno-localisation. (left) representative fields of cells in the indicated conditions. Nuclei borders are indicated by a solid line; (right) quantifications.

### VLP-delivered RnhA rescues the progression of replication forks under stress

Having established that VLP-delivered RnhA is enriched in the vicinity of replication forks, we asked whether it can rescue the progression of replication forks in response to stress. We first incubated asynchronous U2OS cells with VLPs carrying either mCherry, RnhA or *d*RnhA. After 3.5 hours of incubation, we did consecutive pulses with the thymidine analogues IdU and CldU. During the CldU pulse, cells were exposed or not to 50 µM Hydroxyurea (HU) to induce fork slowdown ^27,28^. As expected, in cells pre-treated with VLPs carrying the control protein mCherry, CldU tracks were significantly shorter than IdU tracks in the presence of HU (CldU/IdU ratio ∼ 0.6) (Fig. 3A). Strikingly, cells pre-treated with RnhA-containing VLPs were completely resistant to HU (CldU/IdU ratio ∼ 1). This rescue required the catalytic activity of RnhA because cells pre-treated with *d*RnhA-containing VLPs behaved like cells treated with mCherry-containing VLPs (Fig. 3A). The rescue effect of the over-expression of human RNase H1 on replication fork progression in response to 50 µM HU was shown to require Primpol activity ^27^. Here we show that Primpol is also required for the HU-resistant progression of replication forks in U2OS cells treated with RnhA-containing VLPs (Fig. 3B). Taken together, these observations confirm that VLP-delivered RnhA is active in cells and promotes the Primpol-dependent progression of replication forks in response to low doses of HU in the same way as human RNase H1. A similar strategy was used to show that RnhA-containing VLPs could also rescue the replication fork slowdown associated with loss of Topoisomerase I (Top1) in HeLa cells ^29,30^ (Fig. S3).

**Figure 3:**
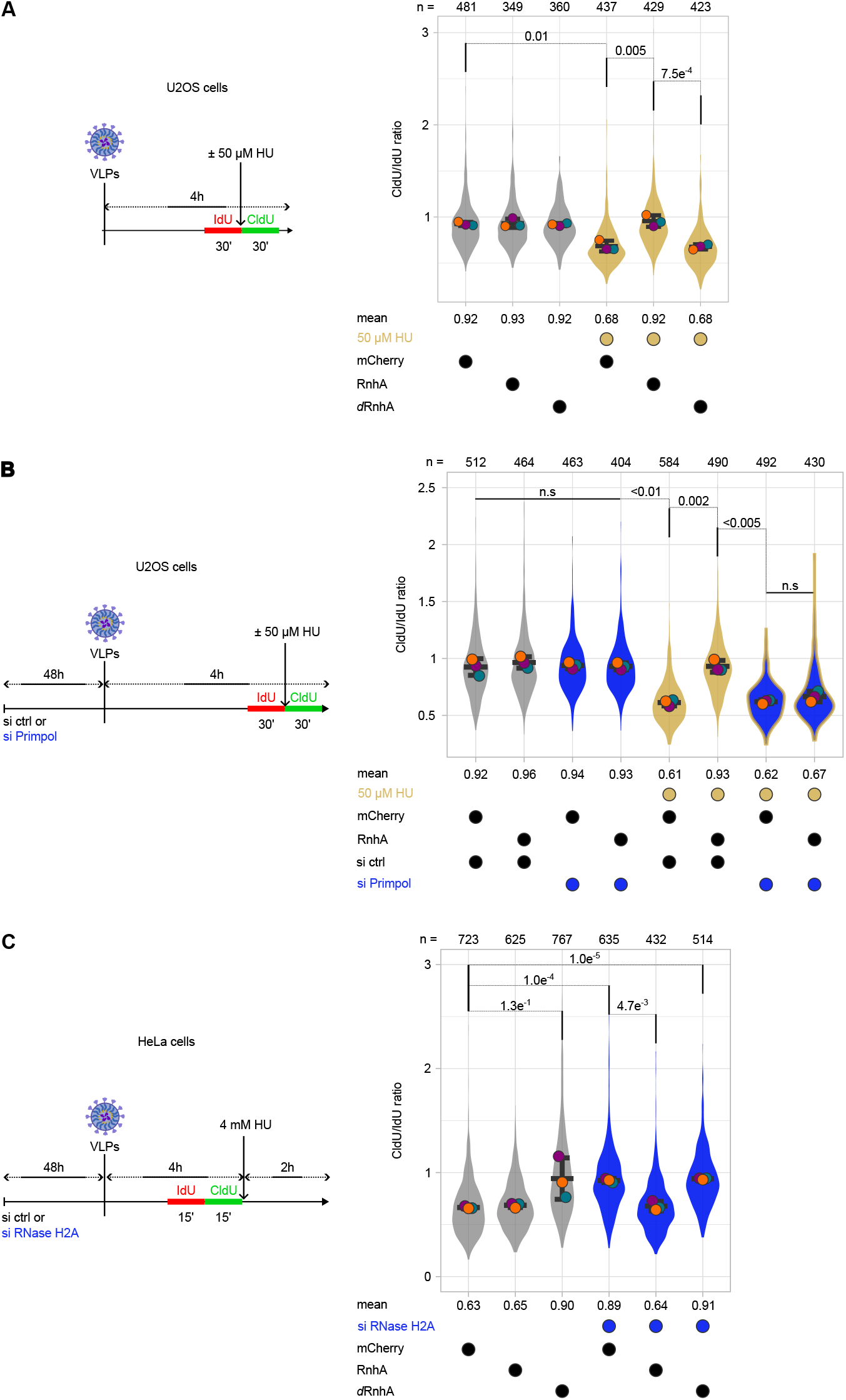
VLP-delivered RnhA rescues the progression and stability of replication forks under stress. A schematic representation of the experiment is shown for each panel. After short pulses with IdU and CldU, chromatin spreading was used to monitor replication fork progression. The impact of VLPs delivering the indicated proteins was assessed **(AB)** in cells treated with 50 µM HU for 30 mins or **(C)** in cells depleted or not of RNase H2A and treated or not with 4 mM HU. The total number of chromatin fibres that were scored is indicated on top of graphs. Welch’s t-test was performed to calculate p-values.

We demonstrated previously that loss of RNase H2A interferes with the resection of stalled replication forks in response to high doses of HU ^14^. We show here that VLP-delivered RnhA could rescue this defect (Fig. 3C). Interestingly, the delivery of the inactive *d*RnhA mutant had a dominant-negative effect on fork resection (Fig. 3C). These observations are consistent with the idea that active RnhA could remove resection-impairing DRIFs, whilst the inactive *d*RnhA might stabilise them. This model is in agreement with the fact that PLA signals with PCNA were stronger for *d*RnhA than for RnhA (Fig. 2). Taken together, these results demonstrate that VLP-delivered RnhA is active in human cells and modulates in *-cis* the behaviour of replication forks in response to a diverse array of replication stresses.

### VLP-delivered RnhA has no significant impact on R-loops mapped by DRIP or Cut&Tag

A common assumption is that the over-expression of RNase H1 promotes the progression of replication forks under stress by removing cotranscriptional R-loops. To test whether VLP-delivered RnhA could remove R-loops genome-wide, we first used the S9.6 antibody to measure the global amount of RNA:DNA hybrids in genomic DNA (gDNA) blotted on a membrane. To increase the signal and assess whether RnhA could remove excessive R-loops, R-loop formation was strongly induced by a short treatment with high doses of Camptothecin (CPT), as described previously ^31^. Surprisingly, S9.6 signals were largely unaffected by the delivery of RnhA, whether or not cells were treated with CPT, whilst they were fully sensitive to an *in vitro* RNase H treatment (Fig. 4A). We confirmed this result using DNA:RNA IP (DRIP-qPCR) at canonical R-loop-forming loci (Fig. S4A) and using both high-resolution, strand-specific, calibrated DRIP-seq (Fig. 4BC) and R-loop Cut&Tag (Fig. S4BC). To test whether VLP-delivered RnhA could remove the DRIP signals induced by the *de novo* transcription of an R-loop forming gene, we used a cell line in which five copies of the R-loop forming *mAirn* gene were integrated under the control of a Dox-inducible promoter ^32^. The addition of Dox for 24 hours induced a ∼10-fold increase in DRIP signals at *mAirn* but this increase also occurred when RnhA-containing VLPs were present during the whole duration of induction (Fig. 4D). This shows that VLP-delivered RnhA could not antagonise the increase in DRIP signals associated with *mAIRN* transcription, even after a 24-hour treatment.

**Figure 4:**
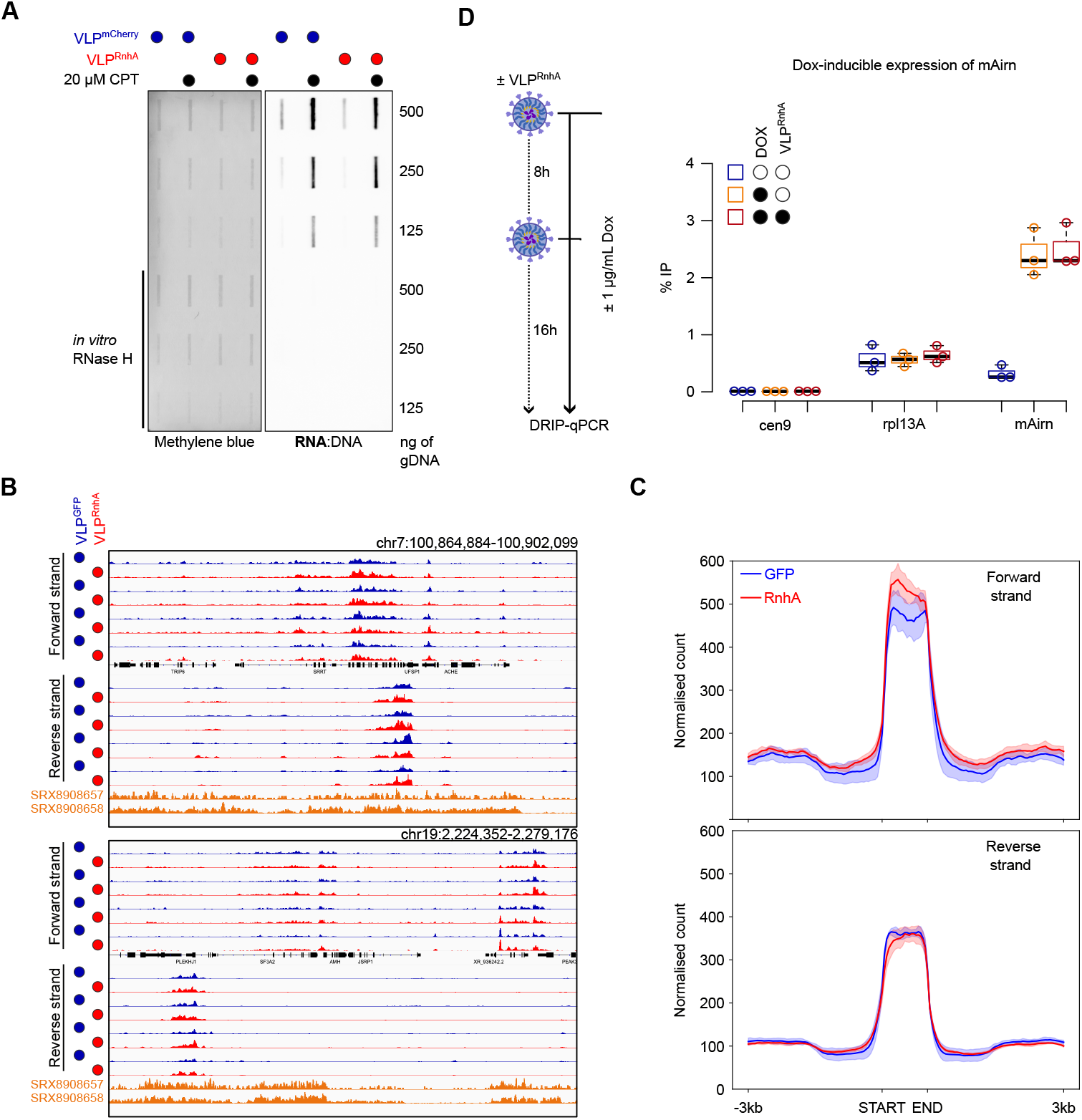
VLP-delivered RnhA does not impact DRIP signals. A. Representative slot blot analysis of total RNA:DNA hybrid levels upon a 4-hour treatment with mCherry- or RnhA-containing VLPs, after cells were treated or not with 20 µM CPT for 5 mins. Methylene blue was used to label DNA. **B**. Examples of calibrated, strand-specific DRIP-seq signals upon treatment with GFP-containing or RnhA-containing VLPs. Published DRIP-seq tracks (SRX8908657&SRX8908658)^35^ are shown for comparison. **C**. Calibrated DRIP signals around called peaks. **D**. qPCR was used to quantify DRIP signals at *mAirn* and at two control loci (cen9 as negative control and Rpl13A as positive control) upon VLP-mediated delivery of RnhA over 24 hours.

We wondered whether the classic over-expression of human RNase H1 (hRNH1) would be able to remove R-loops. A 24-hour over-expression of hRNH1 had no impact on the S9.6 signals measured by slot blot, even after R-loop formation was strongly induced by a short treatment with high doses of CPT (Fig. S5A). These observations were confirmed by DRIP-qPCR (Fig. S5B). Similarly, the transient expression of hRNH1 did not antagonise the increase in DRIP signals associated with *mAIRN* transcription (Fig. S5C). Taken together, our data demonstrate that high levels of active RNase H1 in live nuclei affect the progression of replication forks under stress (Fig. 3) but do not affect overall R-loop levels, as measured by DRIP or Cut&Tag (Fig. 4).

## DISCUSSION

We have shown that VLPs could be used to deliver a controllable amount of ready-made sensors and regulators of RNA:DNA hybrids in live human cells, with unprecedented speed and efficiency and negligible effects on the steady state transcriptome. This approach does not require any genetic modification of target cells and the delivery in parallel of a similar amount of control proteins is easily carried out. This makes this approach particularly attractive to study the role of RNA:DNA hybrids in primary cells. We call this strategy ‘*iCoRD’* for ‘In <u>C</u>ellul<u>o R</u>Nase H <u>D</u>elivery’.

### An under-appreciated RNA:DNA landscape around stressed replication forks

The widely accepted experimental definition of an R-loop is that it is a structure sensitive to an *in vitro* RNase H treatment that maps to a transcribed DNA strand with the same polarity as transcription. If this definition is correct, the striking observation reported here is that the impact of RNase H1 on fork progression and on R-loop resolution can be functionally separated. Consistent with this, we found that the localisation of *d*RnhA in the vicinity of PCNA was resistant to triptolide (Fig. 2), suggesting that fork-proximal RnhA primarily recognizes RNAP2-independent RNA:DNA hybrids. As mentioned above, we propose to collectively call these fork-proximal hybrids ‘DRIFs’, for ‘<u>D</u>NA:<u>R</u>NA hybrids with Impact on <u>F</u>orks’. Our results demonstrate that our VLP-based strategy is able to manipulate the levels of DRIFs in live cells, at least in the stress conditions tested here. We anticipate that DRIFs might have different origins and locations around forks in different genomic contexts and/or in response to different genotoxic treatment. For example, it is likely that VLP-delivered RnhA degrades the recently described, resection-interfering, fork-associated RNA-DNA hybrids (RF-RDs) ^14–16^ that might be stabilised by the loss of RNase H2A in response to high doses of HU (Fig. 3C). It is however unclear at this stage whether transcription-dependent RF-RDs would also form in response to milder replication stresses such as 50 µM HU or the loss of Top1 that we also studied here (Fig. 3 and S3). Importantly, observations made previously in fission yeast suggested that transcription-independent but primase-dependent RNA molecules embedded in reversed forks could prevent the untimely degradation of nascent DNA strands and promote timely replication resumption ^33^, confirming that multiple types of RNA:DNA hybrids distinct from R-loops could impact the behaviour of replication forks upon stress. Taken together, these observations demonstrate the existence of a complex, multi-origin and under-appreciated RNA:DNA landscape around stressed replication forks that regulate the resection of nascent strands upon stress. We anticipate that the VLP-mediated delivery of DRIFs sensors such as *d*RnhA will help in the future to characterise this complex landscape.

### DRIP and Cut&Tag signals are resistant to high levels of RNase H1 in the nucleus

The reason why a large excess of RnhA or hRNH1 in live nuclei could not suppress DRIP or Cut&Tag signals, even when induced *de novo* at the R-loop forming *mAirn* gene, remains unclear. We consider three possible explanations: either (i) the excess of RNase H1 only access a small fraction of RNA:DNA hybrids in cells (such as DRIFs) or, (ii) the rate of R-loop formation is too great for the amount of RNase H1 in nuclei or finally, (iii) DRIP and Cut&Tag do not quantify genuine R-loops. We note that early reports suggested that recombinogenic, transcription-dependent and RNA-containing structures called ‘R-loops’ could form artefactually in test tubes after the extraction of DNA from cells ^34^. Even if those early reports did not formally demonstrate that those were genuine RNase H-sensitive R-loops, similar mechanisms might be at play to explain DRIP signals. Whatever the explanation, an important conclusion of our work is that an increase in DRIP or Cut&Tag signals, even if suppressed by an *in vitro* treatment with RNase H, can only convincingly explain an RNase H1-sensitive phenotype if observed in conjunction with signal suppression by increased cellular RNase H1 levels. As such, we believe that our results invite new interpretations of well-established observations and a change of paradigm to explain the negative impact of transcription on replication.

### Limitations of the study

We cannot be sure at this stage that VLP-delivered RnhA or *d*RnhA could respectively degrade or stabilise all the different types of DRIFs, especially those that might be produced in response to other types of stress than the ones we have tested here. We also do not know whether RnhA is enriched in proximity to all active forks or only the stressed ones.

## ACKNOWLEDGMENTS

We are extremely grateful to Aurèle Piazza and Benoit Palancade for their comments on the manuscript. We thank Jana Dobrovolná and Pavel Janscak for providing the U2OS cell line over-expressing hRNH1 fused to GFP and to Els Verhoeyen for the gift of pBaEVRless. This study was supported by funding from Fondation pour la Recherche Médicale – FRM attributed to AB (contrat doctoral #ECO202306017384), from Ligue contre le Cancer (comité du Rhône) attributed to VV and EPR (Appel d’offres régional 2021), from Fondation ARC pour la recherche sur le cancer attributed to VV (ARCPJA2021060003952), from Agence Nationale de la Recherche (ANR) attributed to PP and VV (project 19-CE12-0016-04 and project ANR-23-CE12-0020-03), from Labex Ecofect (ANR-11-LABX-0048) of the Université de Lyon, within the program Investissements d’Avenir (ANR-11-IDEX-0007) from Agence Nationale de la Recherche (ANR) attributed to EPR and from the European Research Council (ERC-StG-LS6-805500) attributed to EPR under the European Union’s Horizon 2020 research and innovation programs. We acknowledge the contribution of the LBMC Bioinformatics Hub and in particular the help of Laurent Modolo with the analysis of DRIP data. We acknowledge the contribution of SFR Biosciences (Université Claude Bernard Lyon 1, CNRS UAR3444, Inserm US8, ENS de Lyon) and the help of the staff of LyMIC-PLATIM, especially Elodie Chatre, for assistance with confocal microscopy and of Véronique Barateau for assistance with flow cytometry.

## AUTHOR CONTRIBUTIONS

Conceptualisation: VV, EPR; Investigation: AB, VV, Y-LL, CK, YB, VD, LV, LG, SH; Data curation: DC, VV; Formal analysis: DC, VV, AB; Funding acquisition: VV, EPR, PP, PB; Supervision: VV, EPR; Visualisation: AB, DC, VV; Writing-original draft: VV; Writing-review&editing: all authors.

## RESOURCE AVAILABILITY

### Lead contact

Further information and requests for resources and reagents should be directed to and will be fulfilled by the lead contact, Vincent Vanoosthuyse.

### Materials availability

Plasmids generated in this study are available upon request.

### Data and code availability

- The raw Illumina paired-end sequencing data and the processed files in a bigwig format have been deposited to NCBI Gene Expression Omnibus (GEO) under accession GSE288898, GSE288899, GSE288900 and are publicly available from the date of publication.
- All original code has been deposited to Github as described in the methods section and are publicly available from the date of publication.

## DECLARATION OF INTERESTS

The authors declare no competing interests.

## SUPPLEMENTAL INFORMATION TITLES AND LEGENDS

Figures S1-S5

## SUPPLEMENTARY FIGURE LEGENDS

**Figure S1:**
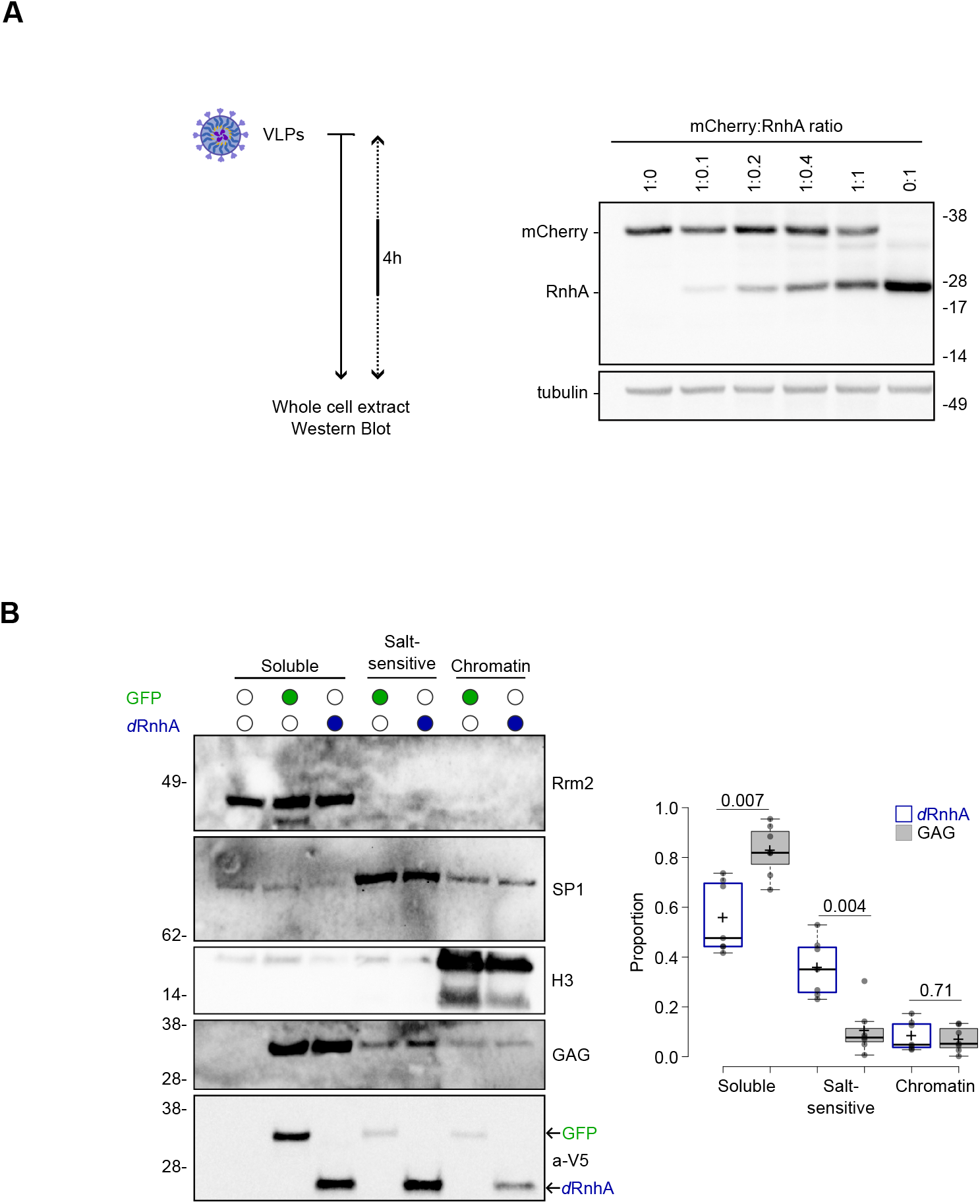
VLP-delivered RnhA associates with chromatin. A. Western blot analysis of total protein extracts of HeLa cells treated with VLPs co-delivering different amounts of V5-tagged mCherry and RnhA. Tubulin was used as a loading control. **B**. HeLa cells were incubated with VLPs delivering either GFP or RnhA^D10N^ (*d*RnhA) for four hours. Protein extracts were fractionated into three fractions: (i) soluble, (ii) salt-sensitive association with chromatin and (iii) tight association with chromatin. Rrm2, SP1 and Histone H3 were used as markers for the different fractions. The retroviral protein GAG and GFP remained mainly in the soluble fraction, whilst a fraction of *d*RnhA associated with chromatin in a salt-sensitive manner. (left) a typical experiment; (right) quantifications of 7 experiments (p-value determined from a Wilcoxon-Mann-Whitney test).

**Figure S2:**
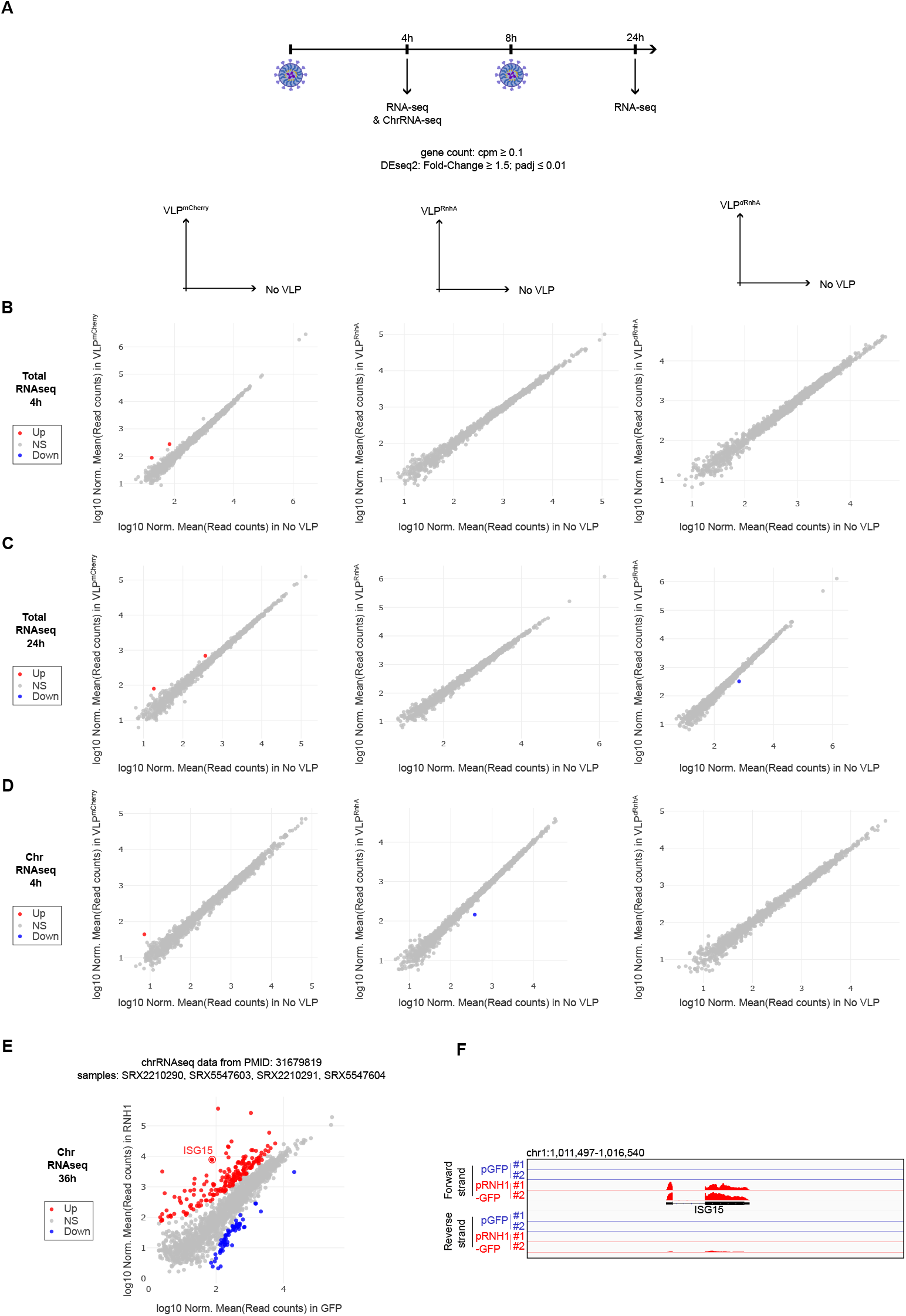
VLP-delivered RnhA does not impact the steady state transcriptome. A. Schematic representation of the experiment. **BCD**. Scatterplot comparisons of the transcriptomes of HeLa cells treated or not with VLPs delivering mCherry (left), RnhA (center) or *d*RnhA (right). Total RNAs were analysed after a 4-hour (**B**.) or a 24-hour (**C**.) incubation. In **D**., chromatin-associated RNAs were analysed after a 4-hour incubation. **EF**. Re-analysis of the indicated published RNA-seq data ^23^ comparing the chromatin-associated RNAs of HeLa cells over-expressing GFP or GFP fused to human RNase H1 for 36 hours. **E**. Scatterplot comparisons. ISG15 is indicated on the plot. **F**. RNA-seq signals around ISG15.

**Figure S3:**
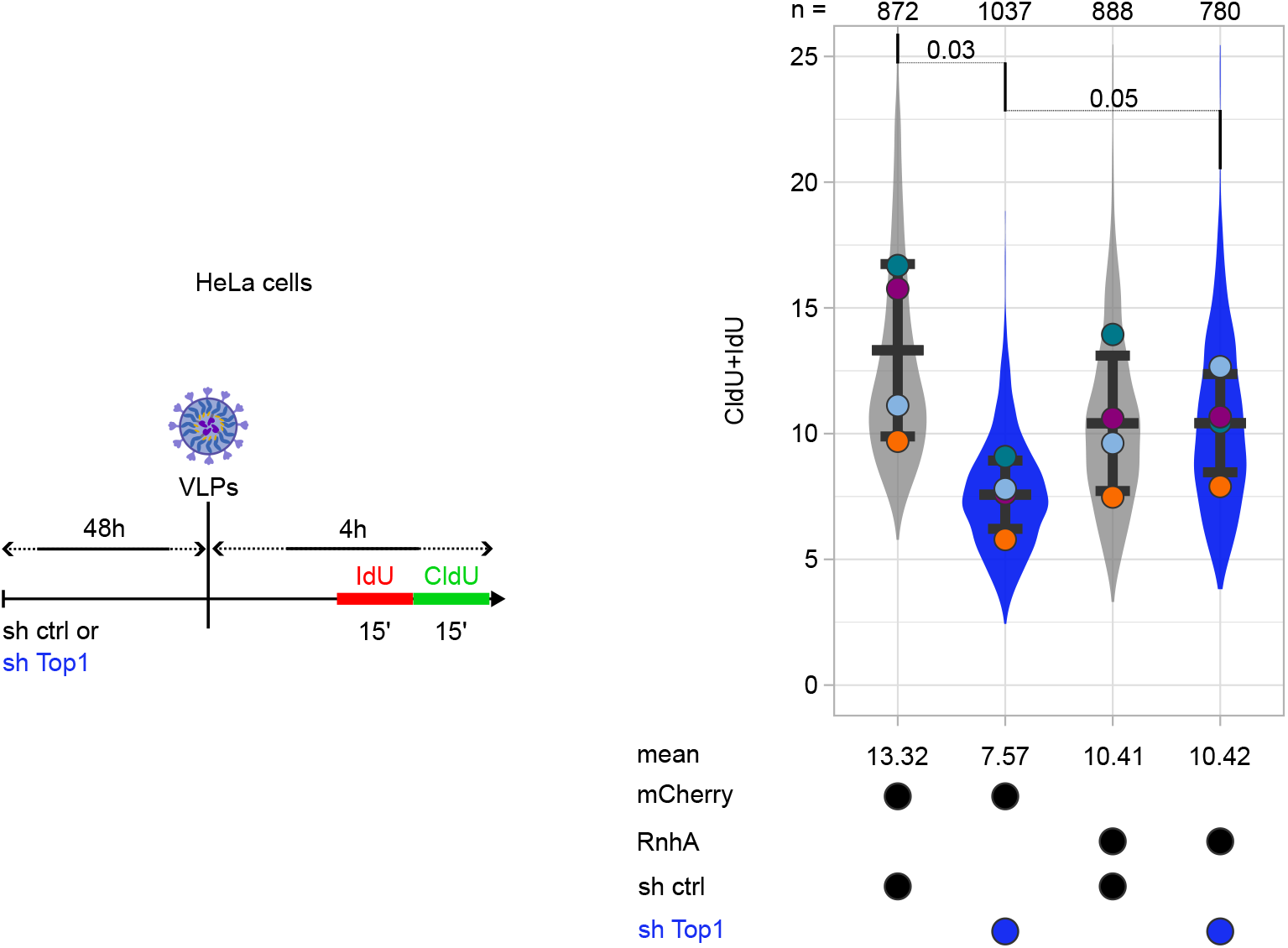
VLP-delivered RnhA rescues the progression of replication forks upon depletion of Top1. A schematic representation of the experiment is shown. After short pulses with IdU and CldU, chromatin spreading was used to monitor replication fork progression. The total number of chromatin fibres that were scored is indicated on top of graphs. Welch’s t-test was performed to calculate p-values.

**Figure S4:**
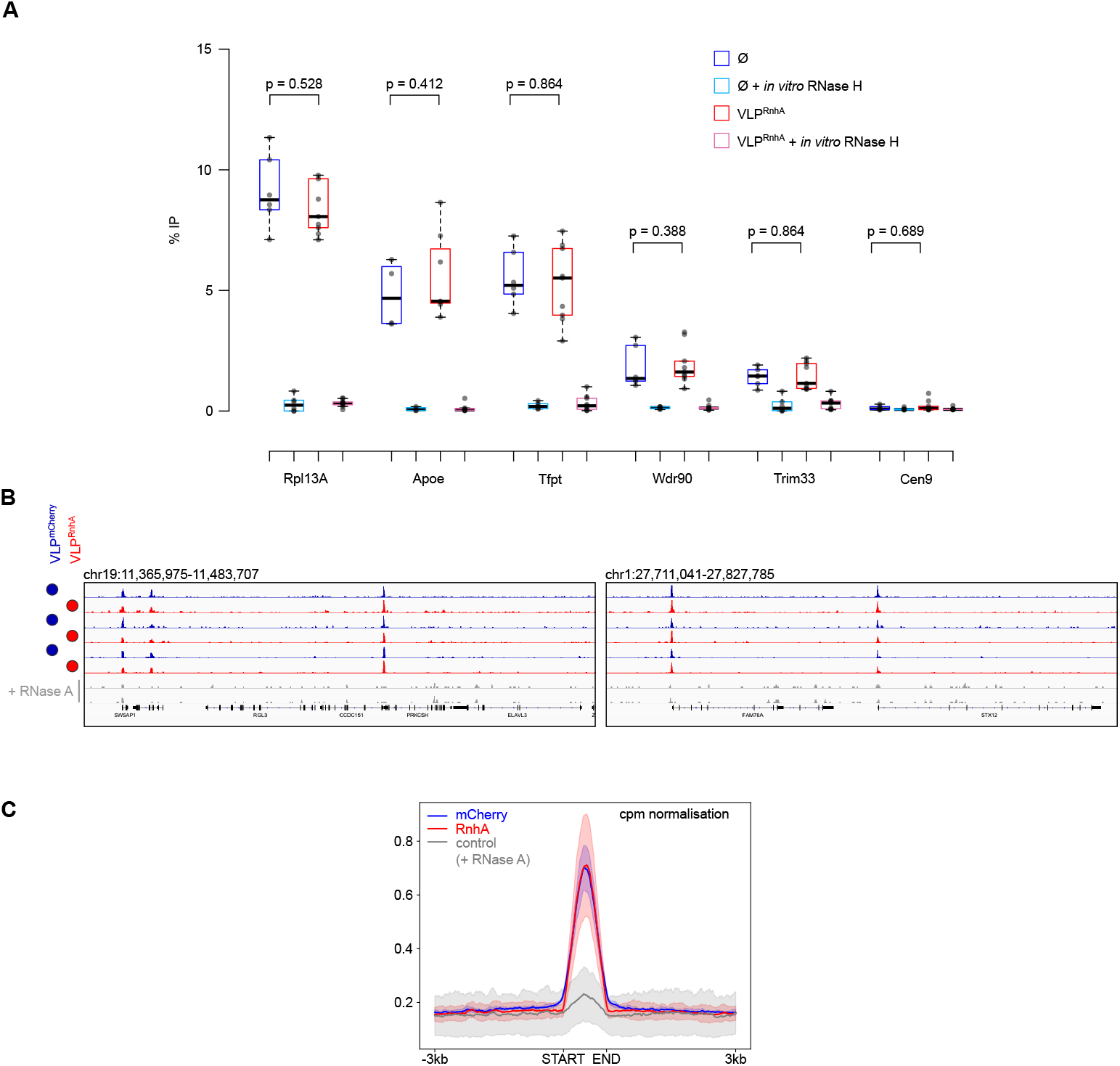
VLP-delivered RnhA does not impact DRIP or Cut&Tag signals. A. DRIP-qPCR signals at different loci in HeLa cells incubated or not (Ø) with RnhA-containing VLPs for four hours. **BC**. Cut&Tag with the S9.6 antibody was carried out to map R-loops after HeLa cells were incubated with either GFP-containing or RnhA-containing VLPs for four hours (n = 3). **B**. Examples of Cut&Tag signals. **C**. Cpm-normalised signals around called peaks.

**Figure S5:**
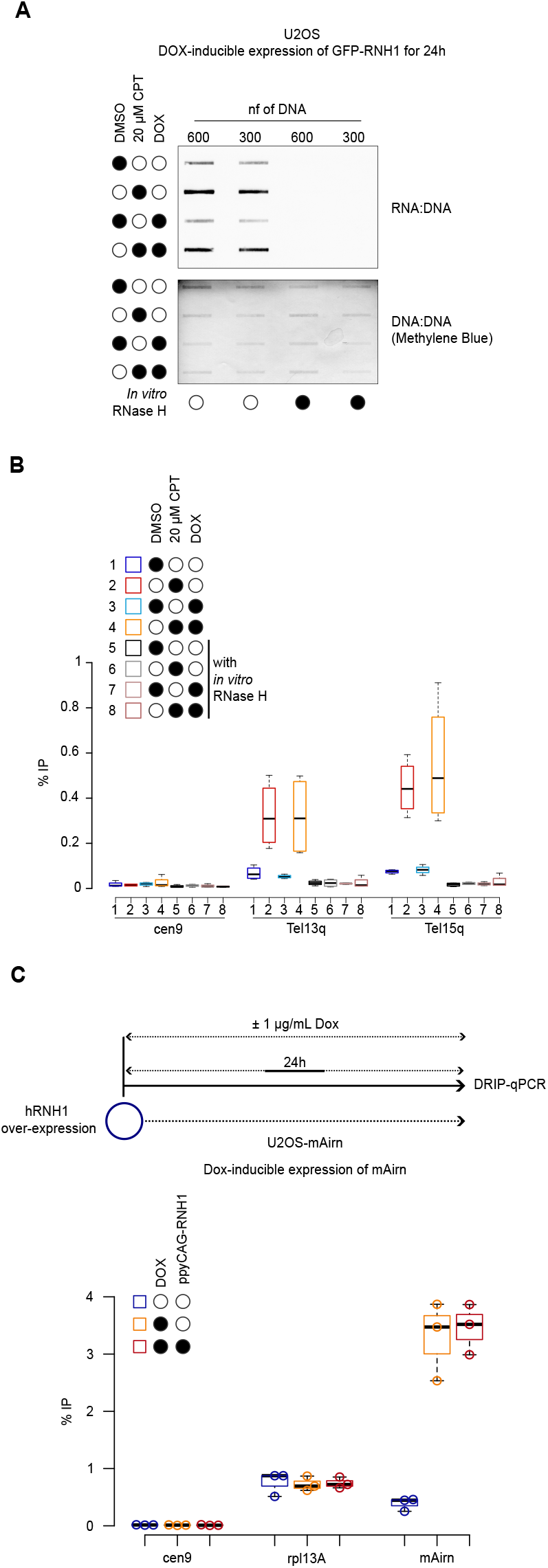
The over-expression of human RNase H1 (hRNH1) does not impact DRIP signals. AB. The over-expression of GFP-hRNH1 was induced or not in U2OS cells by addition of 100 ng/mL of doxycycline for 24 hours. Cells were then treated for 5 minutes with 20 µM CPT or DMSO as control. RNA:DNA hybrid levels were quantified either by slot blotting (**A**.) or by DRIP-qPCR (**B**.) (n = 3). **C**. qPCR was used to quantify DRIP signals at *mAirn* and at two control loci (cen9 as negative control and Rpl13A as positive control) upon over-expression of V5-tagged hRNH1 over 24 hours. 1 µg/mL Doxycycline was used to induce the expression of *mAirn*.

## METHODS

### Cell culture and drug treatment

HEK293T, HeLa and U2OS cells were grown in DMEM medium supplemented with 10% (v/v) foetal calf serum and 1% (v/v) of Penicillin-Streptomycin (Gibco, cat. 15140-122) at 37°C with 5% CO_2_. 1µg/mL puromycin (Gibco, cat. A11138-03) was added to the medium to amplify U2OS cells expressing *mAirn*, as described previously ^32^. 1µg/mL puromycin and 50 µg/mL hygromycin (Gibco, cat. 10687010) were added to the medium to amplify U2OS cells expressing GFP-RNase H1 as described previously ^27^. (S)-(+)-Camptothecin (Merck, cat. C9911) was added at 20 µM final concentration in the growing medium and incubated for 5 minutes. DMSO (Invitrogen, cat. D12345) was used as control. Hydroxyurea (Merck, cat. H8627) was dissolved in the growing medium before being used at the final concentrations of either 50 µM (Fig. 3AB) or 4 mM (Fig. 3C). *mAirn* expression was induced with 1 µg/mL doxycycline for 24 hours (Fig. 4D and Fig S5C). GFP-RNase H1 expression was induced with 0.1 µg/mL doxycycline for 24 hours (Fig. S5AB), as described previously ^27^.

### Protein depletion

Top1 and RNase H2A were depleted as previously described, in ^30^ and in ^14^ respectively. Primpol was depleted using the siRNA GAGGAAACCGUUGUCCUCAGUGUAUUU (Horizon).

### RNH1 over-expression

∼4×10^5^ cells were transfected with 1µg of plasmid ppyCAG-RH1 (Addgene #111906) using jetOPTIMUS (Polyplus, cat. 101000025) according to the manufacturer’s instructions, and incubated for 24 hours. The over-expression of GFP-RNH1 (Fig. S5) was carried out as described previously ^27^.

### Plasmid Construction

Plasmids encoding fusion proteins between Gag (MLV) and the protein of interest were derived from the BIC-Gag-Cas9 plasmid (Addgene plasmid #119942; http://n2t.net/addgene:119942; RRID:Addgene_119942). To generate these constructs, the donor plasmid was digested with AgeI and KpnI to excise the Cas9 coding sequence. The resulting linearised plasmid backbone was purified via gel extraction. Inserts corresponding to the coding sequence of each protein of interest were amplified by PCR, incorporating homology arms complementary to the AgeI- and KpnI-digested vector termini. Assembly of the recombinant plasmids was performed using the NEBuilder HiFi DNA Assembly kit (NEB, cat. E5520S), following the manufacturer’s instructions. pBS-CMV-gagpol was a gift from Patrick Salmon (Addgene plasmid # 35614 ; http://n2t.net/addgene:35614 ; RRID:Addgene_35614). pCMV-VSV-G was a gift from Bob Weinberg (Addgene plasmid # 8454 ; http://n2t.net/addgene:8454 ; RRID:Addgene_8454) ^36^. pBaEVRless was a gift from Els Verhoeyen ^37^.

### VLP preparation and transduction

VLPs were prepared as described previously ^20,38^ with minor modifications. Plasmids were transfected into HEK293T cells plated at 3.5 × 10^6^ cells per 10 cm plate 24 h before transfection with the JetPrime reagent (Polyplus) according to the manufacturer’s instructions. 1 µg of the plasmid carrying the fusion between GagMLV and the protein of interest (), 2.7 µg of pBS-CMV-gagpol, 0.3 µg of pCMV-VSV-G and 0.35 µg of pBaEVRless ^37^ were co-transfected. Supernatants were collected after 40-48h and VLPs were purified and concentrated as previously described ^20,38^. Purified VLPs were incubated at 10 µL/mL in growing medium containing 10 µg/mL of polybrene (Merck, cat. TR-1003-G). For longer incubations, the growing medium was replaced with fresh growing medium containing new VLPs after 8 hours.

### Protein extracts and Western blot

Cell pellets were resuspended in 1X Sample buffer at a ratio of 1.2×10^6^ cells/mL. 1X sample buffer was prepared by diluting 4X Bolt LDS Sample Buffer (Invitrogen, cat. B0007), 10X Bolt LDS Sample Reducing Agent (Invitrogen, cat. B0009) and cOmplete, EDTA-free protease inhibitor cocktail tablets (Roche, cat. 05056489001) in ultrapure water. 20 µL of samples were run on Bolt 4-12 %, Bis-Tris Plus WedgeWell gels (Invitrogen, cat. NW04122BOX) and proteins were transferred on 0.2 µm nitrocellulose blotting membrane (Cytiva, cat. 10600015). Membranes were blocked with 5% milk diluted in PBS containing 0.02% Tween20. After incubation with the appropriate antibodies, signals were revealed using SuperSignal West Femto Maximum Sensitivity Substrate (thermoScientific, cat. 34096). Fractionation of protein extracts was carried out using the Qproteome Nuclear Protein Kit (Qiagen, cat. 37582), according to the manufacturer’s instructions. The primary antibodies used were: SV5-Pk1 (BIO-RAD, cat. MCA1360), anti-Rrm2 (Proteintech, cat. 11661-1-AP), anti-histone H3 (Abcam, cat. ab1791), anti-GAG (Abcam, cat.130757), anti-SP1 (Proteintech, cat. 21962-1-AP). The anti-tubulin was purified from TAT-1 cells (RRID:CVCL_T980)^39^. The HRP-coupled secondary antibodies used were: Mouse IgG HRP Linked Whole Ab - Cytiva NA931-1ML and Rabbit IgG HRP Linked Whole Ab - Cytiva NA934-1ML.

### Immuno-precipitation & slot blot

5-6×10^6^ HEK293T cells were incubated for 4 hours with VLPs carrying *d*RnhA. Cells were washed twice with cold phosphate-buffered saline (PBS), scraped in cold PBS and incubated in hypotonic buffer (20 mM Tris-HCl pH 7.4; 15 mM NaCl; 0.1% Triton X-100; 0.5% IGEPAL) for 15 minutes on ice, in the presence of protease inhibitors (cOmplete EDTA-free protease inhibitor cocktail, Roche, cat. 05056489001). The chromatin fraction was pelleted (5min, 800g), washed in hypotonic buffer, pelleted again (5min, 2000g) and solubilised for 10 min on ice in Solubilisation buffer (10 mM Tris-HCl pH 7.4; 200 mM NaCl; 0.2% Sodium deoxycholate; 0.1% SDS; 0.5% Triton X-100) in the presence of protease inhibitors. Chromatin was then sonicated to ∼600 bp fragments using a Pixul apparatus (Active Motif) with the following parameters: Pulse (50N), PRF (1 kHz), Burst rate (20 Hz), time (15min). After centrifugation (5min, 6000g), 10% of the supernatant were kept for Western blotting (INPUT sample). The rest was diluted in Dilution buffer (10 mM Tris-HCl pH 7.4; 200 mM NaCl; 0.5%Triton X-100) supplemented with protease inhibitors. *d*RnhA was captured on 50 µL V5-Trap® Magnetic Particles (Proteintech, cat. v5td) prepared according to the manufacturer’s instructions. After a one-hour incubation at 4°C on a rotating wheel, magnetic beads were collected on a magnetic rack and washed four times in Wash buffer (10 mM Tris-HCl pH 7.4; 150 mM NaCl; 0.05% IGEPAL; 0.5 mM EDTA). 10% of the beads were kept for Western blotting (IP sample). Nucleic acids were eluted from the remaining beads by a 30-minute incubation in Elution buffer (SDS 1%; NaHCO_3_ 0.1 M). Supernatants were collected into fresh tubes and digested with 200 µg of Proteinase K for 1h at 37°C. Nucleic acid were recovered using Phenol:Chloroform:Isoamyl Alcohol [25:24:1] purification and Ethanol precipitation. After extensive washes with fresh Ethanol 80°, pellets were resuspended in Tris-lowEDTA (10 mM Tris-HCl pH 8.0; 0.5 mM EDTA) and digested with 0.04 U/µL RNase III (NEB, cat. M0245L) in CutSmart Buffer (NEB, cat. B6004S) for 2h at 37°C. For negative controls, the samples were also digested with 0.05 U/µL of RNase H (NEB, cat. M0297L) in the same buffer. The different digests were deposited on a Hybond^™^-N+ membrane (Amersham, cat. RPN203B) using a slot blot apparatus (Bio-Dot^®^ SF Microfiltration apparatus, BioRad) according to the manufacturer’s instructions. Two successive pulses with UV (0.12J) were used to crosslink nucleic acids to the membrane. After blocking the membrane with 5% milk dissolved 1X PBS containing 0.02% Tween-20, RNA:DNA hybrids and dsDNA were quantified respectively using the S9.6 antibody (lab stock) and the ab27156 (Abcam) antibody. Signals were revealed using SuperSignalTM West Femto Maximum Sensitivity Substrate (ThermoScientific, cat. 34096).

### ImmunoLocalisation

∼2.5×10^5^ HeLa cells were plated on coverslips (Marienfield, cat. 0101050). 24h later, cells were treated as indicated, washed twice with cold PBS and pre-extracted with cold PBS-Triton 0.2% (Sigma-Aldrich, cat. T8787) for 2 min on ice. After 2 washes with PBS, cells were fixed in the dark for 15 min with freshly-made PBS containing 3% formaldehyde (thermoScientific, cat. 28908). After 2 washes with PBS, cells were permeabilized with PBS-Triton 0.2% for 5 min and then washed again twice with PBS. Cells were then blocked in a humidity chamber with PBS containing 3% BSA (PBS-BSA) for one hour, and incubated with the relevant primary antibodies in PBS-BSA overnight at 4°C. After 2 washes with PBS-BSA and 1 wash with PBS, coverslips were incubated for 30min in a humidity chamber with the relevant secondary antibodies diluted in PBS-BSA. After 2 washes with PBS, cells were incubated with 0.5 µg/mL DAPI (5min) and coverslips were mounted with VECTASHIELD® Antifade Mounting Medium (Vector Laboratories, cat. H-1000). Image acquisition was carried out using a confocal microscope LSM800 (Zeiss). The following primary antibodies were used: SV5-Pk1 (BIO-RAD, cat. MCA1360) against V5-tagged proteins and anti-RNA-Polymerase II RPB1 Antibody (BioLegend, cat. 920204) against Rpb1. The following secondary antibodies were used: F(ab’)2-Goat anti-Mouse IgG (H+L) Cross-Adsorbed Secondary Antibody, Alexa Fluor™ 568 (Invitrogen, cat. A11019) and F(ab’)2-Goat anti-Rabbit IgG (H+L) Cross-Adsorbed Secondary Antibody, Alexa Fluor™ 488 (Invitrogen, cat. A1070).

### Cell synchronisation

HeLa cells were incubated for 18 hours with 2 mM thymidine (Sigma-Aldrich, cat. T1895). After three washes with the growth medium, cells were released into fresh medium for 200min. When required, 1 µM Triptolide (Sigma-Aldrich, cat. T3652) was added 80min after the release for 120min.

### Flow cytometry

Cells were incubated with 25 µM EdU (Invitrogen, cat. C10632) for 30min. After one wash with PBS, cells were collected, then washed twice with PBS containing 3% BSA (PBS-BSA). To monitor nascent DNA, EdU incorporation was quantified using the Click-iT Plus EdU Flow Cytometry Assay Kit (Invitrogen, cat. C10632) following the manufacturer’s instructions. Total DNA was quantified using 0.5 µg/mL DAPI. Flow cytometry was carried out using a MACSQuant VYB cytometer (Miltenyi Biotec).

### Proximity Ligation Assay (PLA)

∼1.5×10^5^ HeLa cells were used per experiment. Cells were washed twice with cold PBS and then incubated for 10 min with cold PBS containing 0.5% Triton (Sigma-Aldrich, cat. T8787) and protease inhibitors (cOmplete EDTA-free protease inhibitor cocktail, Roche, cat. 05056489001). After 2 washes with PBS containing protease inhibitors, coverslips were fixed for 15 min in the dark with fresh PBS containing 4% formaldehyde (thermoScientific, cat. 28908) and protease inhibitors. After 2 washes with PBS, cells were permeabilized with PBS containing 0.2% Triton for 10 min and then washed twice with PBS. Proximity Ligation Assay was carried out using NaveniFlex Cell MR Red (Navinci NC.MR.100.Red) according to the manufacturer’s instructions. The primary antibodies used were SV5-Pk1 (BIO-RAD, cat. MCA1360) against V5-tagged proteins and anti-PCNA Antibody (Abcam, cat. ab18197) against PCNA. Coverslips were mounted using Duolink® In Situ Mounting Medium with DAPI (Sigma-Aldrich, cat. DUO82040). Image acquisition was carried out using a confocal microscope LSM800 (Zeiss).

### Clustering of PLA signals

All .czi stack files were analysed using the procedure explained here: https://gitbio.ens-lyon.fr/LBMC/RMI2/mundi_centro. Briefly, for every stack, each z slice was saved as an independent .tiff file. Nuclei in each z slice were detected on the DAPI channel using the pre-trained nuclei model of Cellpose ^40,41^. ROIs for each nucleus along the z axis were reassociated using the *AgglomerativeClustering* function of scikit-learn ^42^. For each nucleus, downstream analysis was carried out on the slice with the bigger area. Pixel values and ROIs properties were saved to dedicated files and a dedicated statistical analysis was used to sort nuclei into different clusters depending on the intensity of their fluorescent signals. Briefly, nuclei in all conditions and all biological replicates were pooled and analysed together in an unbiased manner. The distribution of pixel intensities in each nucleus was determined and used to classify nuclei into four different clusters using the kmeans function of scikit-learn ^42^. The proportion of nuclei in each cluster was determined for each condition across biological replicates.

### DNA fibre spreading

DNA fibre spreading was performed as described previously ^43,44^. Briefly, sub-confluent cells were sequentially labelled with 10 µM 5-iodo-2’-deoxyuridine (IdU) and with 40 µM 5-chloro-2’-deoxyuridine (CldU) for the indicated times. About 1000 cells were loaded onto a glass slide (StarFrost) and lysed with spreading buffer (200 mM Tris-HCl pH 7.5, 50 mM EDTA, 0.5% SDS) by gently stirring with a pipette tip. The slides were tilted slightly and the surface tension of the drops was disrupted with a pipette tip. The drops were allowed to run down the slides slowly, then air dried, fixed in methanol/acetic acid 3:1 for 10 minutes, and allowed to dry. Glass slides were processed for immunostaining with mouse anti-BrdU to detect IdU (clone B44, BD Biosciences, Ref: 347580), rat anti-BrdU to detect CldU (clone BU1/75, Abcam, ab6326), mouse anti-ssDNA antibodies and corresponding secondary antibodies conjugated to various Alexa Fluor dyes. Nascent DNA fibres were visualized using immunofluorescence microscopy (Zeiss ApoTome). The acquired DNA fibre images were analysed by using MetaMorph Microscopy Automation and Image Analysis Software (Molecular Devices) and statistical analysis was performed with GraphPad Prism (GraphPad Software). The lengths of at least 150 IdU/CldU tracks were measured per sample.

### Slot blot to quantify RNA:DNA hybrids

After two washes with 1X PBS, 1-2×10^6^ cells grown in 6 cm plates were lysed directly on plates with 1 mL of lysis buffer (10 mM Tris pH 8.0, 1% SDS, 600 µg/mL Proteinase K, 20 mM EDTA pH 8.0). The lysate was then scraped off the plate and transferred into a 1.5 mL tube and incubated at 55°C for 4 hours. Purification with Phenol:Chloroform:Isoamyl Alcohol [25:24:1] pH8.0 was carried out in MaXtract High Density tubes (Qiagen, cat. 129065) and nucleic acids were precipitated with 2.5 volumes of Ethanol. Extensive washes with fresh Ethanol 80° were carried out before nucleic acids were resuspended in 10 mM Tris pH 8.0. The concentrations of genomic DNA (gDNA) were determined by quantitative PCR (qPCR) using primers against Alu elements (Alu F: GTGGCTCACGCCTGTAATC & Alu R: CAGGCTGGAGTGCAGTGG). 4 µg of gDNA was then digested with HindIII-HF (NEB, cat. R3104S) and EcoRI-HF (NEB, cat. R3101S) according to the manufacturer’s instructions. For negative controls, 5U/µg of RNase H (NEB, cat. M0297L) was added to the digest. Different amounts of gDNA were deposited on a Hybond^™^-N+ membrane (Amersham) using a slot blot apparatus (Bio-Dot^®^ SF Microfiltration apparatus, BioRad) according to the manufacturer’s instructions. Two successive pulses with UV (0.12J) were used to crosslink the DNA to the membrane. After blocking the membrane with 5% milk dissolved 1X PBS containing 0.02% Tween-20, RNA:DNA hybrids and dsDNA were quantified respectively using the S9.6 antibody (lab stock) and the ab27156 (Abcam) antibody. Alternatively, dsDNA was revealed with Methylene blue. The concentrations of genomic DNA (gDNA) in the last dilution of gDNA were determined by quantitative PCR (qPCR) using primers against Alu elements (see above) to confirm that an equal amount of gDNA was deposited on a membrane in all samples.

### DRIP

gDNA was extracted and quantified as above. It was then sonicated using a Pixul apparatus (Active Motif) with the following parameters: Pulse (50N), PRF (1 kHz), Burst rate (20 Hz), time (35min). For negative controls, gDNA was digested for at least 5 hours at 37°C with 5U/µg RNase H in the buffer provided by the manufacturer (NEB, cat. M0297L). 4 or 20 µg of sonicated gDNA was used for DRIP-qPCR or DRIP-seq respectively. The S9.6 antibody was coupled to Dynabeads^™^-Protein A (Invitrogen, cat. 10002D) according to the manufacturer’s instructions. 1% of gDNA was set aside (input). The rest of the gDNA and the S9.6 antibody were mixed in cold IP buffer (10 mM Tris pH8.0, NaCl 140 mM, 0.5% Triton, 1X cOmplete EDTA-free protease inhibitor cocktail, (Roche, cat. 05056489001) using a 2:1 ratio (by mass) and rotated for 90 minutes at 4°C. The immunoprecipitated fraction was then washed twice in IP buffer, once with Wash III buffer (10 mM Tris pH 8.0, 1 mM EDTA pH 8.0, 1% IGEPAL, 1% Sodium deoxycholate, 250 mM LiCl) and twice with TE buffer (10 mM Tris, 1 mM EDTA pH 8.0). The immunoprecipitated gDNA fragments were eluted from the beads in Elution buffer (Tris 50 mM, EDTA 10 mM, SDS 1%, 400 µg/mL Proteinase K) at 55°C for 45’ and then purified using Phenol:Chloroform:Isoamyl Alcohol [25:24:1] and Ethanol precipitation. After extensive washes with fresh Ethanol 80°, the eluted fraction was resuspended in 10 mM Tris, 0.5 mM EDTA pH 8.0.

### Strand-specific sequencing libraries

DRIP DNA was treated with 10 µg of RNase A (30’, 37°C) then purified on AMPure beads (1.8x ratio) according to the manufacturer’s instructions. ∼1 ng of DNA was used to prepare sequencing libraries with the xGen™ ssDNA & Low-Input DNA Library Preparation Kit (IDT), according to the manufacturer’s instructions. Sequencing libraries were cleaned twice with AMPure beads (0.85x ratio) according to the manufacturer’s instructions and then quantified using the Qubit DNA high sensitivity kit (Thermo Scientific, cat. Q32851) on a Qubit 2 fluorometer (Thermo Scientific, cat. Q32866). Libraries were paired-end sequenced (2×150 bp) on an Illumina Novaseq6000 platform by Novogene UK.

### DRIP-seq analysis

Paired-end reads were processed, aligned on the GRCh38 genome and calibrated using the strategy described at https://gitbio.ens-lyon.fr/LBMC/Bernard/chipseq/-/tree/dev_accel_1splus?ref_type=heads, using the ‘--keep_multi_map’ option. Reads mapping to the nuclear genome were calibrated over the reads mapping to the mitochondrial genome. Peak calling was carried out with MACS2 with the following options: ‘--bdg --SPMR --nomodel -f BAMPE --min-length 300 --keep-dup all --broad’. Subsequent analyses were carried out with Deeptools on the annotated peaks whose q-value was ≥ 2 and whose length was ≥ 600 bp.

### Cut&Tag

Cut&Tag was carried out with the CUT&Tag-IT® R-loop Assay Kit according to the manufacturer’s instructions (Active Motif, cat. 53167). Libraries were quantified using the Qubit DNA high sensitivity kit (Thermo Scientific, cat. Q32851) on a Qubit 2 fluorometer (Thermo Scientific, cat. Q32866) and paired-end sequenced (2×150 bp) on an Illumina Novaseq6000 platform by Novogene UK. Paired-end reads were processed and aligned on the GRCh38 genome using the nf-core cutandrun 3.1 pipeline (https://nf-co.re/cutandrun/) ^45^ using the cpm normalisation mode.

### RNAseq

Total RNAs were extracted with Trizol (Invitrogen™, cat. 15596026) using the manufacturer’s instructions. To extract chromatin-associated RNAs, cells were first incubated for 15’ at 4°C in Nuclei buffer (Tris 20 mM, NaCl 15 mM, Triton 0.1%, NP-40 0.5%). Nuclei were collected by a 5’ centrifugation at 800 g (4°C). The pellet was washed twice in nuclei buffer and chromatin-associated RNAs were extracted with Trizol (Invitrogen™, cat. 15596026) using the manufacturer’s instructions. For both total RNAs and chromatin-associated RNAs, residual genomic DNA was digested with the TURBO DNA-free™ kit (Invitrogen, cat. AM1907), following the manufacturer’s protocol. RNA quality and concentrations were determined using the Agilent High Sensitivity RNA ScreenTape system, following the manufacturer’s protocol. Sequencing libraries and paired-end sequencing (2×150bp) was performed by Novogene UK. Paired-end reads were processed and aligned on the GRCh38 genome using the nf-core rna-seq 3.9 pipeline (https://nf-co.re/rnaseq/) ^45^. After filtering out genes whose CPM < 0.1, differential analysis with DESeq2 and scatterplot production (padj < 0.01; foldChange 1.5) was carried out using DEBrowser v1.26.3 (http://nasqar2.abudhabi.nyu.edu/DEBrowser/).

## Notes

### Competing Interest Statement

The authors have declared no competing interest.

